# Host association induces genome changes in *Candida albicans* which alters its virulence

**DOI:** 10.1101/2020.04.22.056002

**Authors:** Amanda C. Smith, Meleah A. Hickman

**Affiliations:** Emory University

## Abstract

*Candida albicans* is an opportunistic fungal pathogen of humans that is typically diploid yet, has a highly labile genome that is tolerant of large-scale perturbations including chromosomal aneuploidy and loss-of-heterozygosity events. The ability to rapidly generate genetic variation is crucial for *C. albicans* to adapt to changing or stress environments, like those encountered in the host. Genetic variation occurs via stress-induced mutagenesis or can be generated through its parasexual cycle, which includes mating between diploids or stress-induced mitotic defects to produce tetraploids and non-meiotic ploidy reduction. However, it remains largely unknown how genetic background contributes to *C. albicans* genome instability *in vitro* or *in vivo.* Here, we tested how genetic background, ploidy and host environment impact *C. albicans* genome stability. We found that host association induced both loss-of-heterozygosity events and genome size changes, regardless of genetic background or ploidy. However, the magnitude and types of genome changes varied across *C. albicans* strains. We also assessed whether host-induced genomic changes resulted in any consequences on growth rate and virulence phenotypes and found that many host derived isolates had significant changes compared to their parental strains. Interestingly, host derivatives from diploid *C. albicans* predominantly displayed increased virulence, whereas host derivatives from tetraploid *C. albicans* had mostly reduced virulence. Together, these results are important for understanding how host-induced genomic changes in *C. albicans* alter the relationship between the host and *C. albicans*.

## Introduction

Host-pathogen interactions are multi-faceted. As a fungal opportunistic pathogen of humans, *Candida albicans* has many different relationships with the host. Typically, *C. albicans* is commensal, residing in many niches in the human body, including the gastrointestinal and urogenital tracts, oral cavity, and the skin (1). However, *C. albicans* can be pathogenic and cause superficial mucosal infections and deadly bloodstream infections (2, 3). Much of the research regarding *C. albicans* has focused on its virulence factors, which include, filamentation, biofilm formation, secretory aspartyl proteinases (SAPs), and candidalysin production (4, 5). Host immune cells control *C. albicans* infection by recognizing fungal cells and producing antimicrobial peptides (AMPs) (6) and reactive oxygen species (ROS) which inhibit growth and cause DNA damage (7, 8) However, how host-induced genome alterations in *C. albicans* impact its relationship with the host is not well understood.

Genomic alterations in *C. albicans* have important consequences in other clinical contexts, most importantly in acquiring resistance to the drugs used to treat fungal infections. Analysis of clinical isolates and laboratory studies show that chromosomal aneuploidy and homozygosis of hyper-active resistance alleles are associated with increased antifungal drug resistance (9–14). The ability to generate genetic variation and tolerate genomic perturbations is a strategy that *C. albicans* leverages to adapt to changing environments (15). As a highly heterozygous diploid (16), the *C. albicans* genome is highly labile and undergoes large-scale genome rearrangements like loss-of-heterozygosity (LOH) events and aneuploidy more frequently than small-scale DNA mutations. Furthermore, exposure to physiologically relevant stress conditions, including high temperature, oxidative stress, and antifungal drugs, increases LOH rates even further (17, 18).

Another mechanism *C. albicans* uses to generate genetic variation is parasex, which involves diploid-diploid mating to produce tetraploids (19, 20). Tetraploids undergo stochastic chromosome loss to return to diploid. This process is termed ‘concerted chromosome loss’ and results in reassortment of alleles, loss of heterozygosity (LOH), and aneuploidy (21–23). Tetraploid LOH rate is substantially higher than in diploids (23). In the context of antifungal drugs, large-scale mutation rates are disproportionately elevated in tetraploids compared to diploids (18). Tetraploids, which undergo more genomic changes than diploids, have high potential to produce phenotypic changes with fitness consequences (23, 24). In fact, diploids will produce transient tetraploids via mitotic errors when exposed to high doses of the antifungal drug fluconazole (25). Therefore, tetraploidy may be a useful evolutionary strategy to produce genetic variation.

Clinical *C. albicans* tetraploids have been isolated from human hosts (24, 26–28), although whether they arise via parasex or stress-induced mitotic defects is not known. Furthermore, clinical diploid isolates also carry genomic changes such as aneuploidy and LOH (29). A small number of experimental studies have investigated *C. albicans* genome stability within host environments and found that host-association elevates mutation rates compared to *in vitro* (30– 32). Host-induced genetic variation increases phenotypic variation including colony morphology, hyphal formation, and virulence (30, 32–34). For example, chromosome 6 and chromosome 7 trisomies arise in *C. albicans* associated with murine hosts and display attenuated virulence (31, 33).

Here we investigated how host-association impacts *C. albicans* genome instability across multiple genetic backgrounds and ploidies. We used three diploid-tetraploid pairs of *C. albicans* strains from distinct genetic backgrounds to infect *C. elegans* hosts and extract from to measure LOH frequency and genome size changes. We found that host-association increased genome instability for all *C. albicans* strains, but the degree to which it was elevated depended on strain background. Furthermore, host-associated diploids had minor, but significant genome size changes, whereas host-associated tetraploids rapidly underwent major reductions in genome size. We assessed how these genomic changes altered strain fitness and virulence. Most diploid isolates were more virulent following host-association, but many tetraploid isolates did not change virulence were less virulent, despite undergoing massive genome size changes. Taken together, our results show that host-association induced genetic variation in diploid and tetraploid *C. albicans* of diverse genetic backgrounds which impacted virulence phenotypes.

## Materials and Methods

### Strains and Maintenance

We used six C. albicans strains for this study that vary in their ploidy and their genetic background (Table S1). Each strain was initially struck from glycerol stocks stored at −80°C onto a YPD [yeast peptone dextrose (1% yeast extract, 2% bactopeptone, 2% glucose, 1.5% agar, 0.004% adenine, 0.008% uridine)] plate. After 48 hours at 30°C, a single colony was arbitrarily chosen and considered the “parent strain”.

*C. elegans* N2 bristol (wildtype) were used for fecundity and genome stability assays. *C. elegans* populations were maintained on petri plates containing nematode growth media (NGM) with *E. coli* (OP50) for a food source. *C. elegans* were transferred to a new plate containing freshly seeded *E. coli* every three to four days. For genome stability assays, treatment NGM plates were seeded with both *C. albicans* and *E. coli* and supplemented with 0.2 g/L streptomycin to inhibit overgrowth of *E. coli.* For fecundity and genome stability assays, NGM was supplemented with 0.08g/L of uridine and 0.08g/L of histidine to facilitate growth of auxotrophic *C. albicans* strains.

### Strain construction and verification

To construct diploid clinical strains heterozygous for the *GAL1* locus, we transformed one copy of the *GAL1* open reading frame in FH1 and PN2 with the dominant drug-resistant NAT gene by lithium acetate transformation. *NAT* was amplified from plasmid pMG2120 (35) by PCR (1. 94°C – 5 m, 2. 94°C – 30 s 3. 55°C – 45 s 4. 72°C – 4 m 5. Cycle to (2) 29x 6. 72°C – 10 min) using primers oMH112 and oMH113 listed in (Table 2). Transformants were selected on YPD containing 50 μg/mL NAT. PCR (1.94°C – 3 m. 2. 94°C for 30 s. 3. 55°C – 30 s. 4. 68°C – 2 m. 5. Cycle to (2) for 35x 6. 68°C −10 m.) was performed to verify that NAT was properly integrated using primers oMH106, oMH5, and oMH104 (Table S2). All strains were stored at −80°C and maintained on YPD at 30°C.

### Seeding NGM plates for genome stability assays

Single colonies of *C. albicans* were inoculated into 3 mL of YPD and incubated at 30°C overnight. *C. albicans* cultures were diluted to a final volume of 3.0 OD_600_ per mL in ddH_2_O. Additionally, *E. coli* (OP50) was inoculated into 50 mL of LB and incubated at 30°C for 24-48 hours. Subsequently, *E. coli* (OP50) was pelleted and washed twice with 1 mL of ddH_2_O. The washed pellet was then weighed and diluted to final density of 200mg/mL. *In vitro* treatment plates contained 250 μL of diluted *C. albicans* plated and spread onto an NGM + strep agar plate and incubated overnight at 30°C. *In vivo* treatment plates had 6.25 μL of *C. albicans,* 31.25 μL of *E. coli,* and brought to a final volume of 250 μL with ddH_2_O. This mixture was plated and spread onto an NGM + strep agar plate and incubated overnight at 30°C.

### Seeding NGM plates for fecundity assays

Seeding NGM plates and *C. elegans* population synchronization for fecundity assays were performed as previously described (36). Briefly, *C. albicans* cultures were inoculated into 3 mL of YPD and incubated at 30°C overnight. *C. albicans* cultures were diluted to a final volume of 3.0 OD_600_ per mL. Additionally, *E. coli* (OP50) was inoculated into 50 mL of LB and incubated at 30°C for 24-48 hours. Subsequently, *E. coli* (OP50) was pelleted and washed twice with 1 mL of ddH_2_O. The washed pellet was then weighed and diluted to a final density of 200mg/mL. Day 0 uninfected treatment plates contained 6.25 μL of *E. coli* and brought to a final volume of 50 μL with ddH_2_O. Day 0 *C. albicans* treatment plates had 1.25 μL of C. albicans, 6.25 μL of E. coli, and brought to a final volume of 50 μL. The entire 50 μL was spotted onto the center of a 35-mm-diameter NGM + strep agar plate, followed by incubation at room temperature overnight before the addition of eggs or transferring nematodes. For days 2-7 of the experiment, *C. albicans* treatment plates contained 0.25 μL of *C. albicans,* 1.25 μL of *E. coli* and 8.5 μL of ddH_2_O. For uninfected treatments, 1.25 μL of *E. coli* was mixed with 8.75 μL of ddH_2_O.The entire 10 μL was spotted onto a 35-mm-diameter NGM + strep agar plate, followed by incubation at room temperature overnight before the transfer of nematodes.

### Egg preparation and synchronization for genome stability and fecundity assays

To synchronize *C. elegans* populations, nematodes and eggs were washed off the plate with an M9 buffer and transferred to a 15 mL conical and pelleted at 1200 rpm for 2 minutes. The pellet was resuspended in a 25% bleach solution, inverted for 2 minutes and subsequently centrifuged for 2 minutes at 1200 rpm. The pellet was washed twice with 3 mL of ddH_2_O and resuspended in 1 mL of ddH_2_O. To determine the concentration of eggs, 10 μL was pipetted onto a concave slide, the eggs were counted under a microscope, and the egg suspension was diluted with M9 to a concentration of ~100 eggs per 100 μL.

### Host-associated yeast extractions

*C. elegans* colonized with *C. albicans* and *E. coli* were washed off the plate with 3mL of M9 worm buffer. This suspension was centrifuged for two minutes at 2000 rpm to pellet the worms. The supernatant was removed and 1 mL of 3% bleach was added to the pellet in order to kill any bacteria or yeast that was washed off the petri plate along with the worm. After three minutes of bleach treatment, the worm suspension was centrifuged for one minute at 12,000 RPM. The supernatant was removed and 1 mL of M9 was added to the *C. elegans* pellet and centrifuged for one minute at 12,000 rpm. This step was repeated two additional times to ensure the bleach was removed. The worm pellet was pipetted out into 100 ul aliquots of buffer and transferred to 0.6 mL clear microtubes for manual disruption with a motorized pestle. After one minute of manual disruption, the worm intestine solution was then diluted accordingly with an M9 buffer. All of the intestinal extracts were plated on YPD + 0.034mg/L chloramphenicol to select against any bacteria.

### GAL1 Loss of Heterozygosity Assay

*In vitro:* Single colonies of *C. albicans* were inoculated in 3 mL YPD, grown overnight at 30°C and subsequently diluted to 3 OD in ddH_2_O. 250 μL was plated and spread onto NGM + strep plates and incubated overnight at 30°C and transferred to 20°C for four days. On day four, yeast cells were washed off with ddH_2_O, harvested by centrifugation, washed once with ddH_2_O, resuspended in 1 mL of ddH_2_O and serially diluted for single colony growth. To determine the total cell viability, 100 μL of 10^-6^ dilution was plated onto YPD and grown for 48 hours at 30°C. To identify cells that lost *GAL1,* 100 μL of 10^-2^ and 10^-3^ dilution was plated onto 2-deoxygalactose (2-DOG; 0.17% yeast nitrogen base without amino acids 0.5% ammonium sulfate, .0004% uridine, .0004% histidine, 0.1% 2-deoxygalacose, 3% glycerol) and CFUs counted following 72 hours incubation at 30°C.

*In vivo:* The approach was very similar as the *in vitro* LOH assay described above, with the following changes. A population of ~100 nematodes were plated on each treatment plate containing both *C. albicans* and *E. coli.* On day four, yeast was extracted as described in the previous section. A dilution of 10^-1^ and 10^-2^ was plated on YPD + chlor to enumerate total growth and undiluted cells were plated on 2-DOG to select for the loss of *GAL1.* Three technical replicates were used for each *C. albicans* strain for both *in vitro* and *in vivo* experiments. At least three biological replicates were used for each genome stability assay.

### In Vitro Passaging

Serial passaging experiments were performed as previously described (23). Briefly, we inoculated 36 single colony isolates for each *C. albicans* strain into 500 ul liquid YPD in 96-deep well culture blocks and incubated them at 30° with shaking. Every 24 hours, 5 ul of culture was diluted in 495 ml fresh YPD (1:100 dilution) and incubated at 30° with shaking. On days 4, 7, 14 and 28 cultures were simultaneously prepared for flow cytometry or for long-term storage in 50% glycerol at −80°. Glycerol stocks were also prepared on days 10, 17, 21 and 24.

### Flow cytometry for ploidy determination

Single yeast colonies extracted from *C. elegans* were inoculated in YPD and incubated overnight at 30°C. Samples were sub-cultured into wells with 495 μL of fresh YPD and grown at 30°C for an additional 6 hours. Cells were subsequently collected by centrifugation (1000 rpm, five minutes) and the supernatant removed and resuspended in 50:50 TE (50 mM Tris, pH 8 and 50 mM EDTA). Cells were fixed by adding 180 ul of 95% ethanol and stored overnight at 4°C. Following fixation, cells were collected by centrifugation, washed with 50:50 TE and treated with 50ul of RNase A [1mg/mL] for one hour at 37°C with shaking. Following RNase treatment, cells were collected by centrifugation, RNAase solution was removed and cells were resuspended with 50ul proteinase K [5 mg/mL] and incubated for 30 minutes at 37°C. Following proteinase K treatment, cells were collected by centrifugation and washed once with 50:50 TE and resuspended in 50ul SybrGreen (1:50 dilution with 50:50 TE; Lonza, CAT#12001-798, 10,000X) and incubated overnight at 4°C. Following SybrGreen straining, cells were collected by centrifugation, SybrGreen was removed and cells were resuspended in 50 μL 50:50 TE. Samples were sonicated to disrupt any cell clumping and subsequently run on a FACScaliber flow cytometer. To calibrate the LSRII and serve as internal controls, the reference diploid (SC5314) and tetraploid strains were used.

### Ploidy Analysis

Flow cytometry data was analyzed using FlowJo, by plotting the FITC-A signal against the cell count. Two peaks were observed, the first representing the G1 mean and the second peak representing the G2 mean, which has double the genome content of the G1 peak and therefore twice the fluorescence. Ploidy values were calculated using the G1 mean and compared to standard diploid and tetraploid control strains.

### Fecundity assays

Approximately, 50 eggs were added to each control and treatment plates. After 48 hours of growth at 20°C, a single L4 nematode (x10 per treatment) was randomly selected and transferred to an uninfected or *C. albicans* treatment plate and incubated at 20°C. Each nematode was transferred to a new plate every 24 hours for five consecutive days. Any eggs laid for each 24-hour interval were incubated for 24 hours at 20°C and the number of viable progenies produced per worm was enumerated.

### Growth Rate Assays

12 single colonies for each parent strain and 7 colonies for each host-associated yeast strain were grown overnight in 450 μL with shaking of YPD in a 96-well block at 30°C with shaking. Cultures were diluted 10-fold with ddH_2_O. 15 μL of diluted culture was inoculated into 135 μL of YPD in a sterile round-bottom 96-well plate and placed on the Cytation microplate reader. OD was measured every 15 minutes for 24 hours at 30°C with shaking. Growth rate was determined using a custom R script that calculates the maximal growth rate in each well as the spline with the highest slope from a loess fit through log-transformed optical density data that reflects the rate of cell doubling (developed by Richard Fitzjohn, as in (37)).

### Statistical Analysis

Statistical analysis was performed using GraphPad Prism 8 software. Data sets were tested for normality using the D’Agnostino & Pearson omnibus normality test. To test for specific difference between the frequency of LOH for host and no host treatments, we used an unpaired, Mann Whitney U test. To determine deviation from ploidy *in vitro,* (diploidy or tetraploidy) we pooled the data for all the diploid or tetraploid strains on day 0 to calculate the mean and standard deviation of the population of replicate lines. Any replicate line that was +/− 1 standard deviation away from the pooled mean of day 0, was considered a deviation. The same calculation was used to determine deviation *in vivo,* however a pooled standard deviation was not performed across strains. To test for differences in growth rate and fecundity between host-associated isolates and their respective ancestral strain we used an unpaired, Mann-Whitney U-test.

## Results

### Host-association elevates LOH events in laboratory and clinical C. albicans isolates

The murine host environment elevates genome instability of the laboratory strain of *C. albicans* compared to *in vitro* (30). Using a novel model system to investigate *C. albicans* genome stability, we wanted to determine whether the host environment elevates genome instability regardless of *C. albicans* genetic background. We used a *C. elegans* infection model and compared the frequency of loss-of-heterozygosity (LOH) at the *GAL1* locus (17) of a laboratory and two clinical diploid strains *in vivo* to the *in vitro* LOH frequency (Figure 1A). If the host environment elevated *C. albicans* genome instability, then the host-associated LOH frequency should increase relative to *in vitro.* For the laboratory (11.5x) and the oral clinical (9.7x) strains, we found that host-associated LOH is approximately ten-fold higher than *in vitro* (Figure 1B; teal and purple). However, for the bloodstream clinical strain, there was no significant difference in LOH frequency between the *in vitro* and host-associated treatments, likely due to the high *in vitro* LOH frequency (Figure 1B; orange). Indeed, there were significant differences among the strain backgrounds for in the *in vitro* (p<0.0001, Kruskal-Wallis test) and host-associated (p=0.0008, Kruskal-Wallis test) treatments, with the bloodstream strain displaying higher instability than the other two strains (Table S1). Our results are consistent with data from murine models (30–33) and support the hypothesis that host environments increase *C. albicans* genome instability. Our work also demonstrates that *C. albicans* genetic background influences its genome instability.

**Figure 1:**
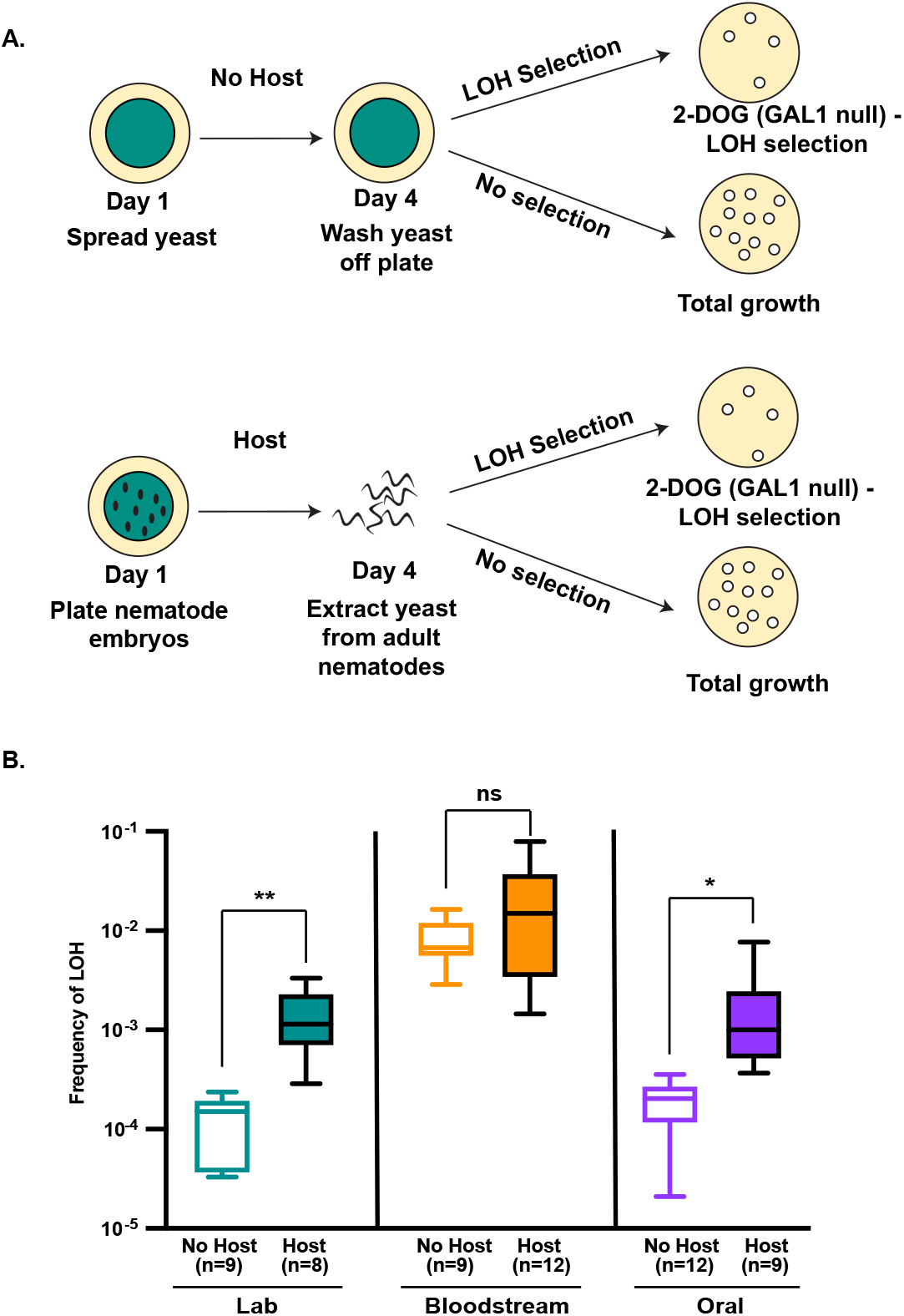
Host-association. induces LOH. A) Experimental schematic. *C. albicans* was grown in the presence with or without nematode hosts. Following four days of exposure, C. albicans were either extracted from host guts or simply washed off the plate. Subsequently, extracted or washed C. albicans were plated on both 2-DOG to select for LOH events of on YPD (rich media) to determine total viable growth. B) Diploid GAL1 LOH rates following no host or host exposure for laboratory (green), bloodstream (orange), and oral (purple) strains. Boxes represent the 25th and 75th quartile with the whiskers representing the minimum data point to the maximum data point. Asterisks indicate any significant differences between the no host and the host treatment groups (* p = .02, ** p = .001; Unpaired Student’s T-

### Host-association induces genome size changes in clinical diploid strains

Sin ce strain background impacts LOH frequency (Fig 1), we next wanted to examine if it also affects genome size changes over time. We have previously shown that clinical strains were more! unstable than the laboratory strain, howeverwe did not investigate changes over the course of s eria l pa ssagi n g . To i d ent ify *in vitro* genome size changes over time, we passaged 60 replicate lines for each genetic background in nutrient-rich liquid media for 28 days and periodically measured genome size via flow cytometry for every replicate line. To assess whether replicate lines deviated in genome size changes over time, we plotted the genome size for each replicate population as the fraction of its initial genome size. Most replicate lines maintained their initial genome size throughout the 28-day experiment, regardless of strain background. However, the number of replicate lines that deviated from diploidy was higher for the clinical genetic backgrounds compared to the laboratory strain (Fig. 2A ‘deviation’). For example, on Day 14, the clinical bloodstream and oral strains had 69% and 72% of their replicate lines with deviations from their initial diploid state. In contrast, only 33% of laboratory replicate lines deviated from diploidy by that same time point. Remarkably, the bloodstream strain had deviations with both major gains (i.e. 1.5x) and losses (i.e. 0.5x) in genome size that suggest that haploidy and triploidy can arise in this strain background (Figure 2A and S1A). For the oral strain, the majority of the deviations in diploidy were minor losses in genome size (Figure S1A). In addition to minor losses in genome size, in the laboratory strain multiple G1 peaks were observed during flow cytometry for several replicate lines on Day 4, indicating mixed populations. However, by Day 28, all of the laboratory replicate lines resolved back to a diploid state (Figure 2A and S1A). Together, these data demonstrate that the two clinical strains are more likely to generate genome size changes than the laboratory strain during *in vitro* passaging.

**Figure 2:**
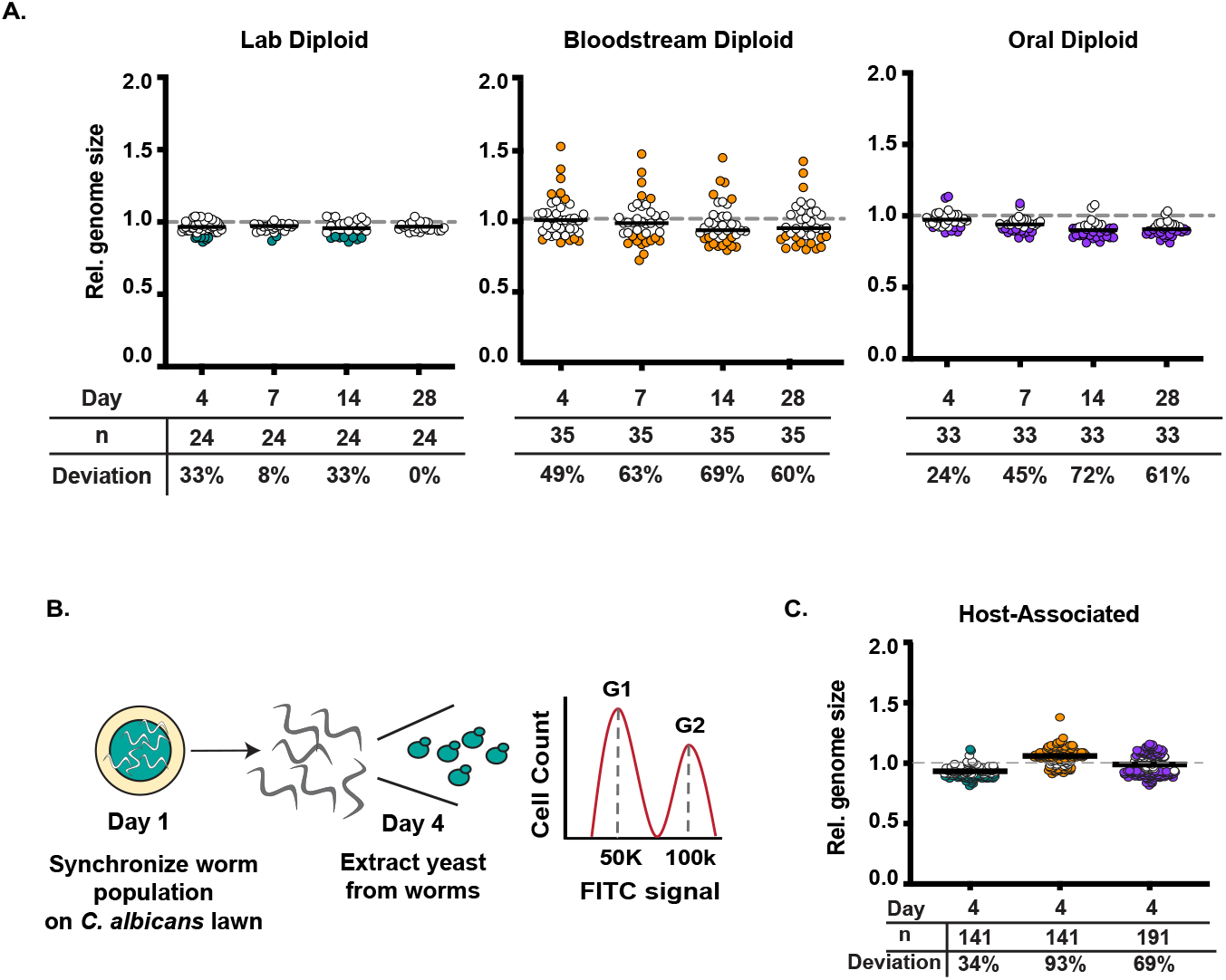
Diploid genome stability *in vitro* and *in vivo*. A) The relative genome size of laboratory, bloodstream, and oral diploid replicate lines passaged for 28 days in rich media was calculated by dividing the genome size (measured via flow cytometry) of each replicate line on each day, by the genome size of day 0 for the same replicate line. Any line that was more than one standard deviation away from tine mean of all lines on Day 0 was considered adeviation from diploidy. Colored circles represent replicate lines that are no longer diploid. B) Experimental salnematic forgenomechanges/n *vivo.* C) After foas days or aseociation with a population of of nematodes, laboratory, bloodstream, and oral isolates were extracted from the gut and genome size was measured via flow cytometry. The relative genome size was calculated by dividing the genome size of each isolate by the mean genome size of the respective *C. albicans* strain not exposed to the host. Deviation from diploidy was considered to be more than 1 SD away from the mean of each strain background. The median was plotted for each of the hostassociated isolates (solid black line). Colored circles represent host-associated isolates that were no longer diploid.

Next, we wanted to assess how the host impacts ploidy stability for each of the three strains. Similar to the LOH assay, we exposed hosts to *C. albicans* for four days and subsequently extracted the yeast on day four. Colonies were picked at random and flow cytometry was performed to assess genome size (Figure 2B). As an *in vitro* control, we plated an equivalent lawn of *C. elegans* in the absence of nematode hosts and. We calculated the relative genome size for the host-associated isolates as a fraction of the genome size of the *in vitro* control not exposed to the host. The laboratory strain had the least number of isolates that deviated from diploid (Figure. 2C, teal). Of the 34% of isolates that were no longer diploid, all had minor losses in genome size (Figure 2C and S1B). In the oral diploid strain, 69% of host-associated isolates were no longer diploid. The deviations from diploid in this strain were both minor gains and losses (Figure 2C, purple and S1B). Shockingly, 93% of the bloodstream host-associated isolates were no longer diploid, after exposure to the host for four days. The majority of deviations from diploidy in this strain background were both major and minor gains in genome size (Figure 2C, Orange and S1B). These results are consistent with the laboratory strain being more stable than the clinical strains, both *in vitro* and *in vivo*. Furthermore, our results indicate that the host environment increases genome instability, both in terms of LOH and genome size changes, yet the amount and direction of these genomic changes depends on strain background.

### Tetraploids undergo rapid genome size reduction in the host environment

Our previous work demonstrated that a majority of tetraploid isolates were no longer tetraploid after 28 days (23, 24). However, it was unclear how quickly each of these backgrounds began to lose chromosomes. Therefore, we wanted to analyze the genome stability of laboratory and clinical tetraploids over time. For the following experiment, we used three tetraploid strains; the laboratory tetraploid strain, which resulted from the mating between two laboratory diploids, a bloodstream clinical tetraploid recovered from a marrow transplant patient, and a vaginal clinical strain recovered from a vaginal infection. To measure genome size changes over time *in vitro,* we passaged 60 replicate lines for each tetraploid genetic background in nutrient-rich liquid media for 28 days and periodically measured genome size via flow cytometry for each replicate line. To assess whether replicate lines deviated in genome size changes over time, we plotted the genome size for each replicate population as the fraction of its initial genome size. Unlike diploids, most tetraploid replicate lines did not maintain their initial genome size, instead most underwent genome reduction to about half their initial genome content (i.e. approximately diploid after passaging), regardless of strain background. By Day 28, 100% of the vaginal and bloodstream replicate lines were no longer tetraploid, and 97% of laboratory replicate lines were not tetraploid (Figure 3A and S1C). However, we detect genome size changes at earlier timepoints for the clinical strains compared to the laboratory strain. By Day 7, more clinical replicate lines (97% and 74% for bloodstream and vaginal, respectively) had a reduction in genome size compared to the laboratory strain in which only 53% of replicate lines had a reduction in genome size. Similar to our previous work (24), we demonstrated that tetraploids reduce to diploidy after serial passaging. Additionally, we demonstrated that the clinical tetraploids undergo a more rapid reduction in genome size compared to the laboratory strain during *in vitro* passaging.

**Figure 3:**
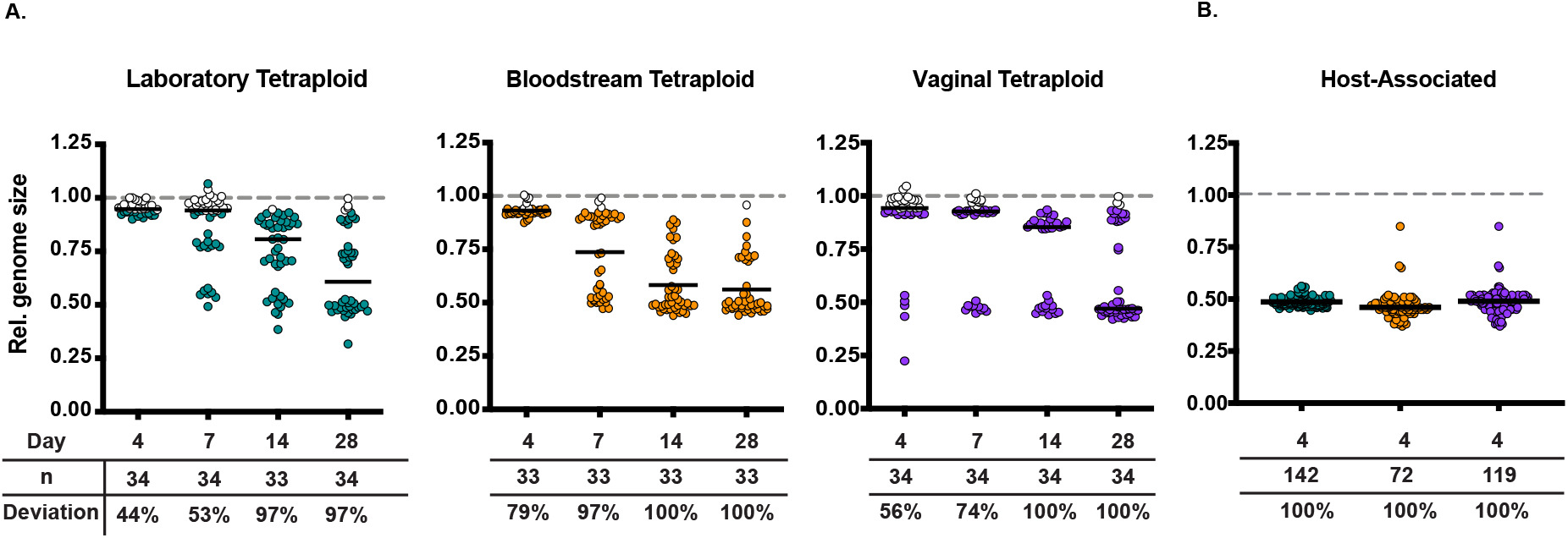
Tetraploid genome *stability in vitro* and *in vivo.* A) The relative genome size of laboratory, bloodstream, and vaginal tetraploid replicate lines passaged for 28 days in rich media was calculated by dividing the genome size (measured via flow cytometry) of each replicate line on each day, by the genome size for Day 0 for the same replicate line. Deviation from diploidy was considered +/− 1 standard deviation away from the mean of all lines on Day 0. Colored circles represent replicates line that are no longer diploid. B) After four days of association with a population ofnematodes,laboratory, bloodstream, and vaginal isolates were extracted from the gut and genome size was measured via flow cytometry. The relative genome size was calculated by dividing the genome size of each isolate by the mean genome size of the respective *C. albicans* strain not exposed to the host. Deviation from tetraploidy was considered to be more than one standard deviation away from the mean of each strain background. The median was plotted for each of the host-associated isolates (solid black line). Colored circles represent host-associated isolates that were no longer tetraploid.

We found that the host induces genome size changes in diploid *C. albicans.* Indeed, many clinical isolates do vary in ploidy (26–28). Therefore, we next wanted to assess whether the host maintains non-diploids. To assess host-associated tetraploid stability, we infected a population of *C. elegans* with each one of the three tetraploid strains mentioned previously. After four days of exposure to *C. albicans,* we extracted *C. albicans* from the *C. elegans* gut via manual grinding and subsequently measured the genome size of the host-associated isolates via flow cytometry. We then calculated the relative genome size compared to the *in vitro* controls for each strain background. After only four days of host-association, 100% of all host-associated single colony isolates were no longer tetraploid, regardless of strain background. Furthermore, the vast majority of these tetraploid-derived isolates were approximately one-half of their initial genome content, and thus, are likely diploid (Figure 3B). This suggests that while the host environment may generate tetraploids, it does not readily maintain tetraploids. Together these data suggest that the host-environment rapidly reduces the genome size of tetraploids.

### Host-induced genome size changes impact C. albicans growth and virulence

Since host-association induced genomic changes in *C. albicans,* we next wanted to assess if these changes had any fitness consequences on growth rate. We focused our attention to host-associated derivatives with genome size changes, since genome size changes were not biased by LOH marker selection and because we did not have host-associated LOH isolates from our three tetraploid strains. For each strain, we selected twelve host-associated isolates (Table S4) and measured their growth rate in nutrient rich media and compared their growth rate to that of their respective parental strain (Fig. 4A & B). It should be noted that there were some differences between genetic background and ploidy state in the parental strains. For each genetic background, the diploid growth rate was significantly different from its corresponding tetraploid growth rate (Table S5). For the laboratory and bloodstream ploidy pairs, the tetraploid grew more slowly than the diploid, but for the oral/vaginal genetic background the opposite was observed. Furthermore, within each ploidy class, the laboratory and bloodstream genetic backgrounds were significantly different from the oral/vaginal genetic background (Table S5). When we compare the host-associated isolates to their respective parental strain, we find collectively that derivatives from all strain backgrounds except the oral diploid and bloodstream tetraploid were significantly different. Furthermore, the diploid genetic backgrounds had decreased growth rates and the tetraploid genetic backgrounds increased growth rates. When we compare the host-derived isolates as individuals to their parental strain, we detect some significant changes in growth for all strain backgrounds and ploidies (Fig. 4 A & B, filled symbols).

**Figure 4:**
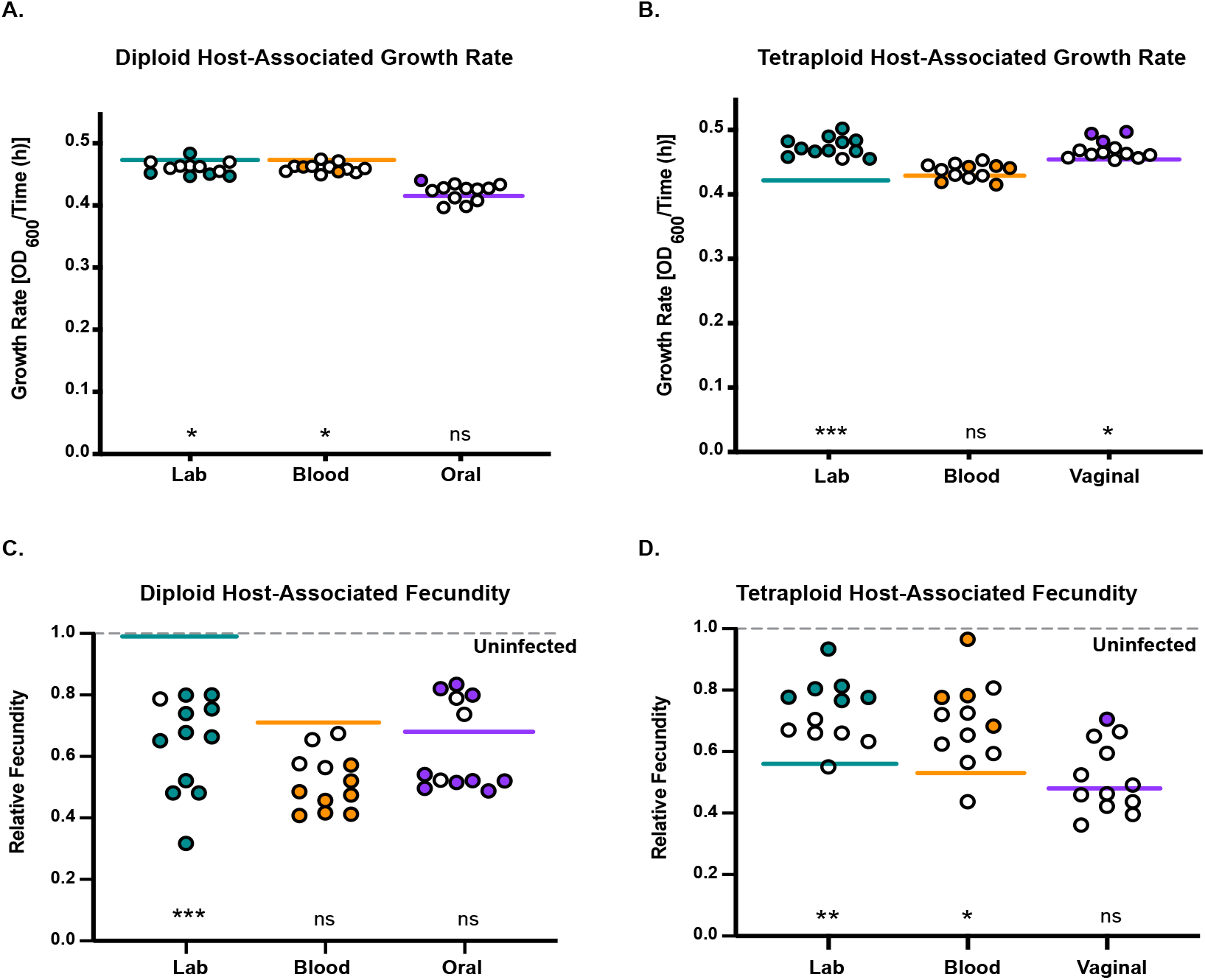
Fitness and virulence consequences associated with host-induced genomic changes. A) Growth rate of the laboratory (green), bloodstream (orange), and oral (purple) host-associated isolates. Each solid line represents the mean growth rate of the ancestral strain. The filled in circles represent host-associated growth rates that are significantly different from the mean growth rate of the ancestral strain (Mann-Whitney U). B) Same analysis as A) instead with tetraploid host-associated isolates C) Relative fecundity of the laboratory (green), bloodstream (orange), and oral (purple) host-associated isolates. Relative fecundity was calculated by dividing the total brood size of nematodes infected with *C. albicans* by the total brood size ot nematodes intected with *E. coli* on*ly (*uninfected; grey dashed line). Solid lines represent the mean relative decundity of each ancestral strain. Fi lled in circles ate significantly different from the ancestral strain, open cincles are not (Manc-Whitney U). D) Same analysis as C) instead with tetraploid host-associated isolates. We tested for differences between the host-associated isolates and their respective parents using a Mann-Whitney U test (*** p <.0001, ** p < .01 * p < .05, ns = not significant)

Decreases in virulence or changes to commensal-like phenotypes have been previously shown following short- and long-term association with host environments (33, 34) To assess whether host-induced genome size changes altered virulence phenotypes, we measured the reproductive success (i.e. fecundity) of *C. elegans* hosts infected individually with each parental strain and their twelve host-derived isolates and calculated the relative fecundity by dividing the brood size of infected hosts by the brood size of uninfected hosts (Figure 4C & D). Similar to the growth rates, it should be noted that there were some differences between genetic background and ploidy state in the parental strains. The diploid-infected fecundity was significantly higher than its corresponding tetraploid-infected fecundity for the laboratory, bloodstream, and oral/vaginal genetic backgrounds, indicating that these tetraploids are more virulent than diploid (Table S6). Genetic background impacted host fecundity for diploids but not tetraploids (Table S6). When we compare the host-associated isolates to their respective parental strain, we find collectively that derivatives from the laboratory diploid and tetraploid, and the bloodstream tetraploid were significantly different. We found that the laboratory diploid derivatives reduced brood size, therefore, increased in virulence whereas the tetraploid derivatives increased in brood size, and therefore decreased in virulence. When we compare the host-derived isolates as individuals to their parental strain, we detect significant changes in host fecundity for all strain backgrounds and ploidies (Fig. 4 C & D, filled symbols). In general, many of the diploid host-derived isolates were more virulent than their respective parental strain, with the exception of a small number of the host-derived isolates of the vaginal strain background, that displayed reduced virulence. All of the tetraploid host-associated isolates that significantly differed showed reductions in virulence. However, there was no direct correlation between genome size changes and relative fecundity or growth (Figure S2). Taken together, our results indicate that even short periods of host association induce genomic changes that have direct impacts on virulence phenotypes.

## Discussion

Here, we investigated how diverse genetic backgrounds and ploidy states of *C. albicans* impact genome stability both inside and outside of the host environment. Previous work has shown that differences in genetic background give rise to phenotypic differences (29), and also impact long-term genome dynamics (24). Furthermore, earlier studies have shown that the host environment increases genome instability (30–33), but these studies do not address the roles genetic background and ploidy play in host-associated genome dynamics. By using three diploid-tetraploid pairs of *C. albicans* strains with distinct genetic backgrounds we were able to compare strain differences in genome stability both *in vitro* and *in vivo.* Furthermore, we analyzed how host-induced genetic changes impacted growth and virulence phenotypes. We found that host-association increased genome instability relative to *in vitro* for all strain backgrounds (Figure 1). However, the magnitude by which the host elevated genome instability was dependent on strain background. We observed that diploids had minor genome size changes occur in the host environment (Figure 2), whereas our three tetraploid strains underwent rapid and large genome reductions in the host environment (Figure 3). Finally, when assessing whether host-induced genome size changes impacted virulence, we found diploid derivatives generally increased in virulence and tetraploids generally decreased in virulence (Figure 4).

We were surprised to detect significant virulence changes in our isolates derived from diploid-infected hosts, since we only detected modest genome size changes. However, 28 out of the 36 host-associated isolates had significant changes in virulence compared to their parental strain (Figure 4C). Virulence increased in the derivatives from both the laboratory and bloodstream strains, as well as for half of the derivatives from the oral genetic background. Interestingly, 25% (3/12) of the derivatives from the oral strain background had reductions in virulence. Furthermore, the level of diploid genome instability depended on genetic background. For example, host-induced LOH was ~10-fold elevated in the laboratory and oral genetic backgrounds, but only 2-fold increase in the bloodstream strain. This is most likely due to the already elevated LOH frequency of the bloodstream strain *in vitro.* These findings are consistent with whole-genome sequencing studies that show naturally occurring LOH events in clinical isolates (29, 38). These sequencing studies also detected chromosomal aneuploidy in a small number of clinical strains (28, 29). In our work, we observed that clinical diploid strains undergo minor genome size changes more frequently over the course of *in vitro* passaging and host-association compared to the laboratory strain (Figure 2). This study builds upon two important experimental studies which found that host environments elevate genome instability in *C. albicans* (30–32) and we extend this by explicitly testing for differences in multiple genetic backgrounds as well as in tetraploids.

Tetraploids underwent massive genome reductions when exposed to the host environment, regardless of genetic background, and in contrast to diploids, which had significant but modest genome size changes. We propose that the host environment is inherently stressful and drives genome instability in *C. albicans* similar to stress-induced mutagenesis. There are several physiological relevant stressors that elevate LOH rates *in vitro,* including high temperature and oxidative stress (17, 39). Reactive oxygen species (ROS) production is an innate immune defense used to defend the host against invading pathogens (7, 8), which inhibits growth by inducing DNA damage (40). Our results indicate that host-induced genome instability could result from host ROS production. Given that *C. elegans* has a conserved innate immune system that includes producing ROS to defend against pathogens (41–43), it would be interesting to investigate how immune function contributes to pathogen genome instability.

The extreme genome instability observed in tetraploids, is partly due to intrinsic highly labile nature. It has been well established that tetraploid *C. albicans* has higher levels of genome instability compared to diploids (18, 22–24), and this phenomenon is also observed in related yeast species (44–47). However, the impact of genetic background on tetraploid genome stability has been extremely limited to date (24). We have previously shown strain dependent differences in tetraploids following long-term *in vitro* serial passaging (24). In this current work, we detected early differences across genetic backgrounds in tetraploid genome reduction during *in vitro* serial passaging (Figure 3A). From our collective *in vitro* results, we anticipated that host-association would likely induce genome size changes to some degree in tetraploids. However, we were surprised to observe all tetraploid host-derivatives with genome reductions, most of which were close to diploid in content (Figure 3B), after only four days of host-association. While tetraploid *C. albicans* have been isolated in clinical settings (26–28), they are rare in comparison to diploid clinical isolates. Our results showing rapid host-induced tetraploid genome reduction may help explain the rarity of clinical tetraploids.

Host-induced genome changes resulted in subsequent changes in virulence phenotypes for all strains (Figure 4C & D). We did not anticipate much phenotypic variation because host-induced genetic changes arose in the absence of selection, so it was striking that ~78% and ~31 % of diploid and tetraploid host-associated derivatives changed virulence relative to their parental strains. Not only were virulence changes more frequent in the diploid host derivatives, nearly all were increases in virulence. In contrast, the few significantly different tetraploid host derivatives had reductions in virulence. A potential reason for this difference stems from differences in baseline virulence between the parental diploid and tetraploid strains (Table S6), in which tetraploids were generally more virulent than diploids.

Finally, we were surprised that so few of the host-derived tetraploids changed in virulence, given the frequency and magnitude of genome size changes induced in the host environment. There was no correlation between changes in genome size and changes in virulence (Figure S2), similar to previous work demonstrating that *in vitro* adaptation to nutrient depletion is not correlated to genome size changes (24). Tetraploid genome reduction generates massive karyotypic heterogeneity in cell populations through its parasexual cycle (22, 23) by re-assorting chromosomes into new allelic combinations, that has been proposed to facilitate rapid adaptation (15). In our study, we measured genome size changes, but haven’t yet characterized the allelic composition in our host-derived isolates. When we consider the large genomic but small phenotypic changes in tetraploids, coupled with the small genomic but large phenotypic changes in diploids, we propose that specific allelic combinations and potential *de novo* mutations foster virulence changes.

## Acknowledgments/Author Contributions

We would like to thank Judy Dinh for her technical support, Dr. Levi Morran and Ognenka Avramovska for critical reading of the manuscript, and Emory University School of Medicine Flow Cytometry Core. This research is supported by NSF DEB-1943415 and Emory University startup funds awarded to MAH.

ACS and MAH designed the study. ACS constructed the necessary strains and conducted *in vivo* genome stability, LOH, fecundity, and growth rate assays. MAH conducted *in vitro* passaging experiments. ACS analyzed the data. ACS and MAH wrote, reviewed and edited the manuscript.

## SUPPLEMENTAL FIGURES AND TABLES

**Supplemental Figure S1:**
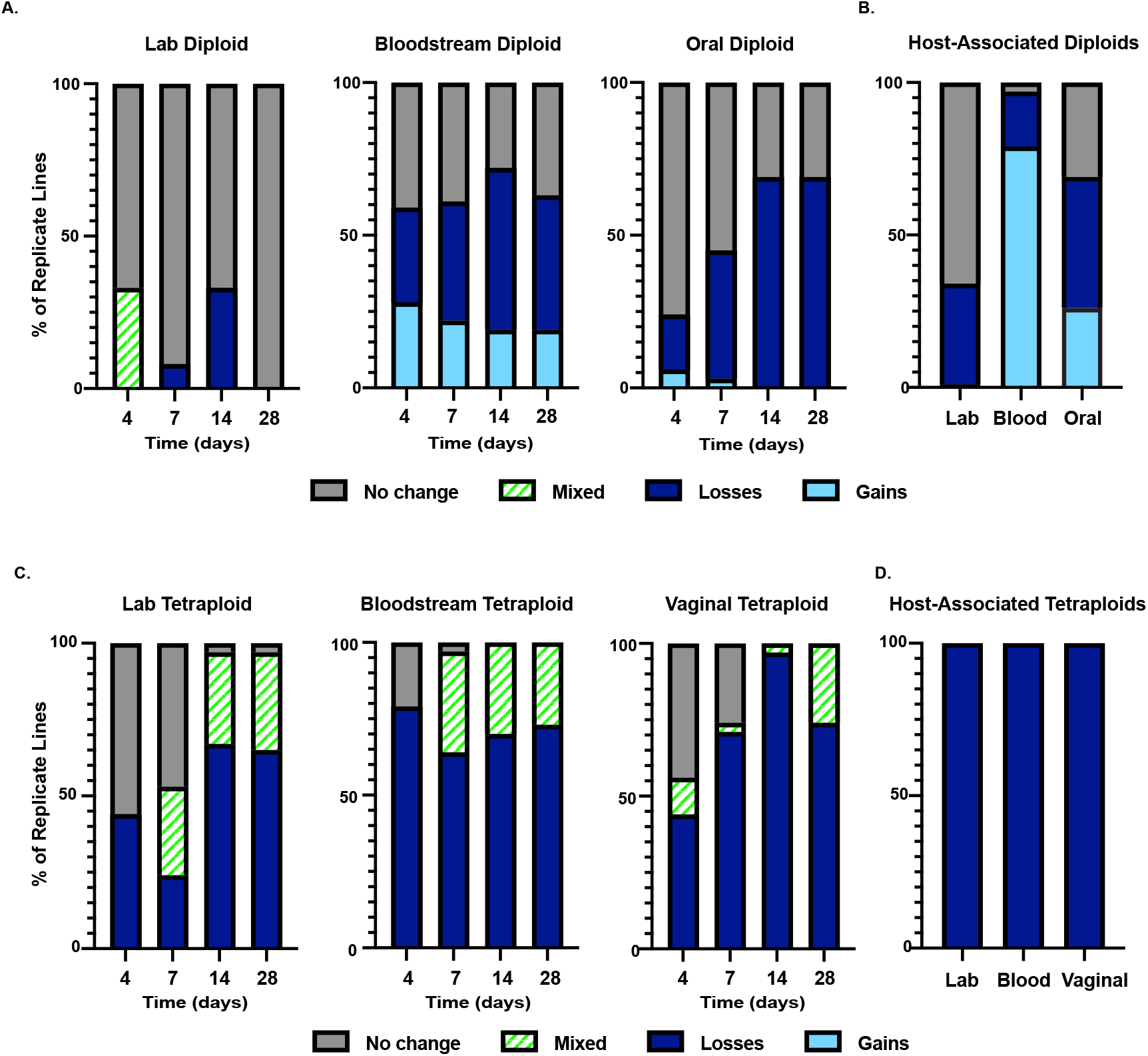
Types of genomic changes observed in diploids and tetraploids. A) The percent of genome size gains and losses in diploid strains. Flow cytometry was used to determine genome size. Anything greater than one standard deviation from the mean of day 0 for all replicate lines was considereda gain in genome size (Light blue). Anything less than one standard deviation fro m the m ean of day 0 for all replicate l i nes was considehed a loss in genome size (dark blue). Any replicate line that had more than one peak when analyzed via flow cytometry was considered a mixed population (green and white). B) Same analysis as A, but for host-associated diploids. C) Same analysis as A, but for each of the tetraploids *in vitro*.

**Supplemental Figure S2:**
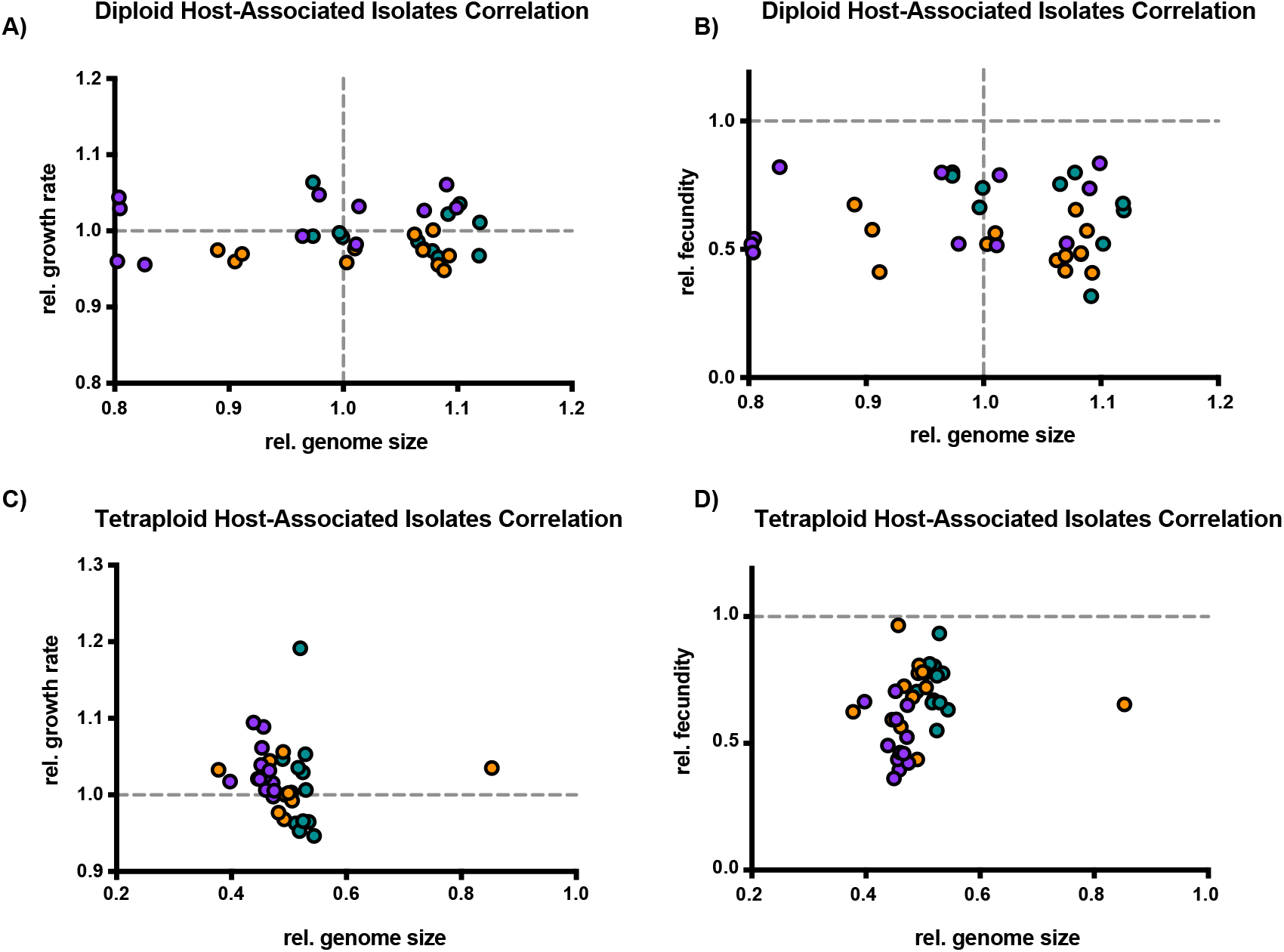
Correlation between genome size changes and fitness or virulence in both tetraploid and diploid *C. albicans* host-associated isolates. A) Correlation between the relative growth rate and the lelative genome size for each of the twelve host-associaied laboratory (green), bloodstream (orange), and oral (purple) isolates. Relative growth rate was calculated by dividing the growth rate of each host-associated isolate by the mean growth rate of the ancestral strain. Relative genome size was calculated by dividing the genome size of each host-associated isolate by the mean genome size ofthe ancestral stcain. (Pearson correlation; laboratory: r = – .08, p = .805 bloodstream: r = .18, p = .57 oral: r = .38, p = .22). B) Correlation between the relative fecundity and the relative genome size for each of the twelve host-associated laboratory (green), bloodstream (orange), and oral (purple) isolates. Relative growth rate was calculated by dividing the growth rate of each host-associated isolate by the mean growth rate of the ancestral strain. Relative fecundity was calculated by dividing the fecundity of each host-associated isolate by the mean fecundity of the uninfected control. (Pearson correlation; laboratory: r = – .57, p = .051 bloodstream: r = −.31, p = .33 oral: r = .42, p = .18). C) Same correlation as A, instead with the tetraploid laboratory (green), bloodstream (orange), and vaginal (purple) host associated isolates. (Pearson correlation; laboratory: r = – .24, p = .45 bloodstream: r = .08, p = .81 vaginal: r = −.23, p = .47). D) Same correlation as B, instead with the tetraploid laboratory (green), bloodstream (orange), and vaginal (purple) host associated isolates. (Pearson correlation; laboratory: r = – .09, p = .78 bloodstream: r = −.04, p = .91 vaginal: r = −.33, p = .29).

**Supplemental Table S1:**
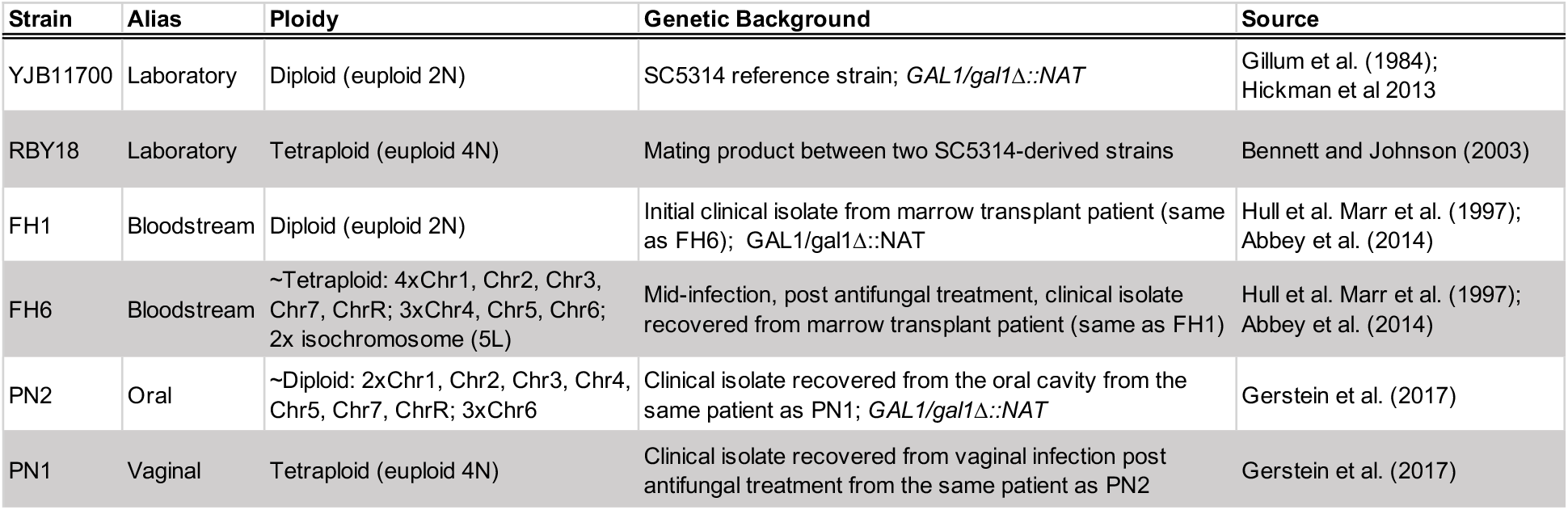
Strains used in this study

**Supplemental Table S2:**
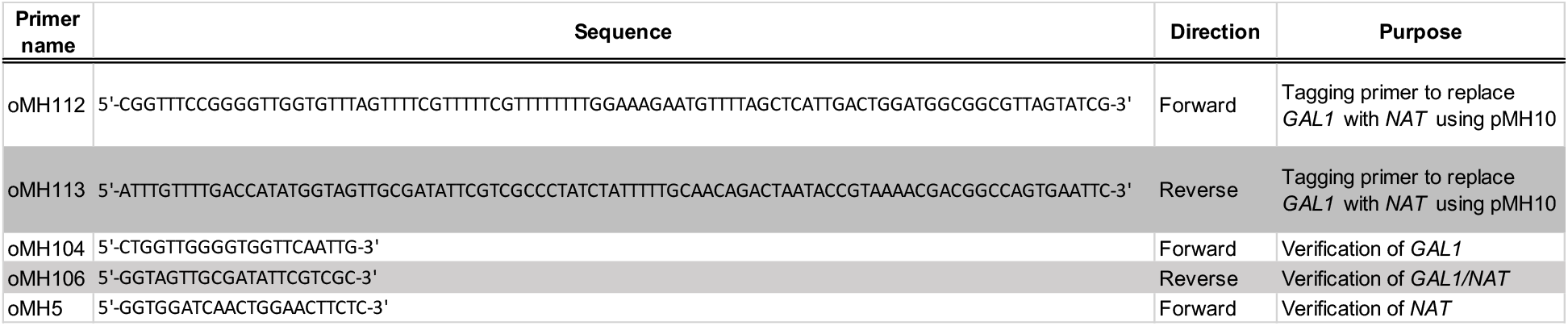
Oligos used in this study to generate *gal1::NAT/GAL1* heterozygous loci

**Supplemental Table S3:**
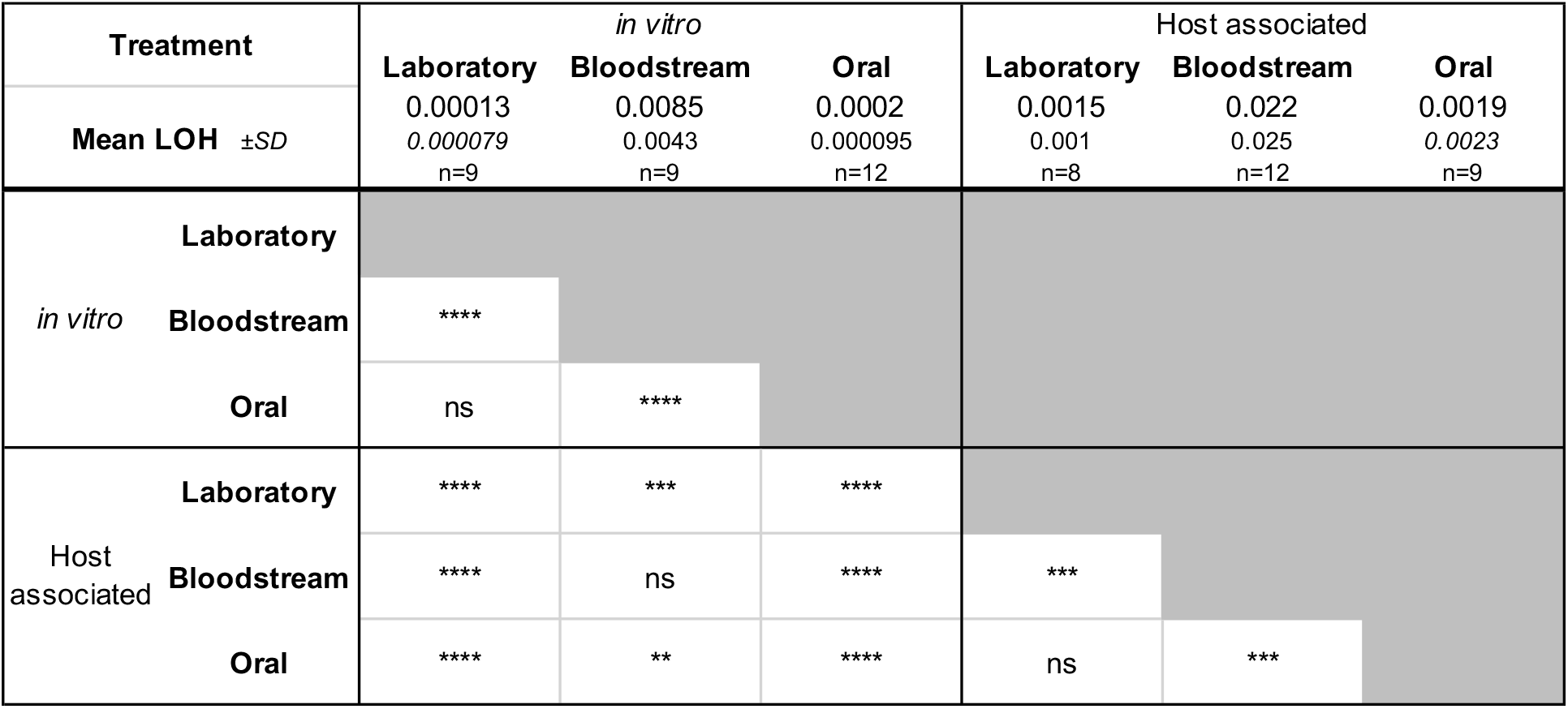
Diploid LOH frequency. Mean LOH frequency, standard deviation (SD) and number of biological replicates (n) are indicated for each treatment. Differences between treatments were tested in all pairwise combinations using the Mann Whitney U test and significance is indicated (ns – not significant; * p<0.05; ** p>0.01; *** p>0.001; **** p>0.0001).

**Supplemental Table S4:**
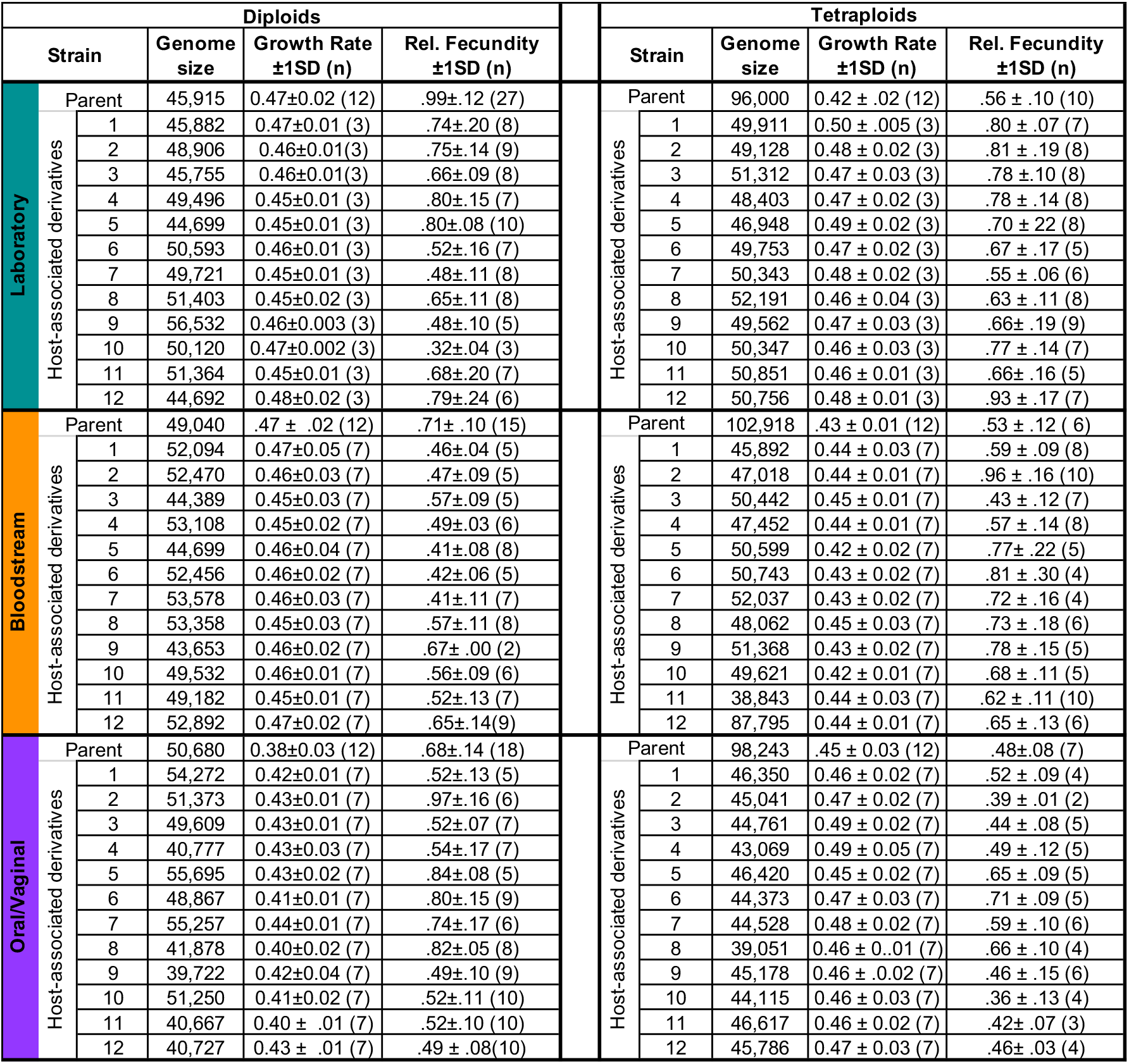
Characteristics of host-associated isolates. For each genetic background and ploidy, the genome size (measured as mean G1 FITC signal), growth rate, and infected host fecundity values are indicated.

**Supplemental Table S5:**
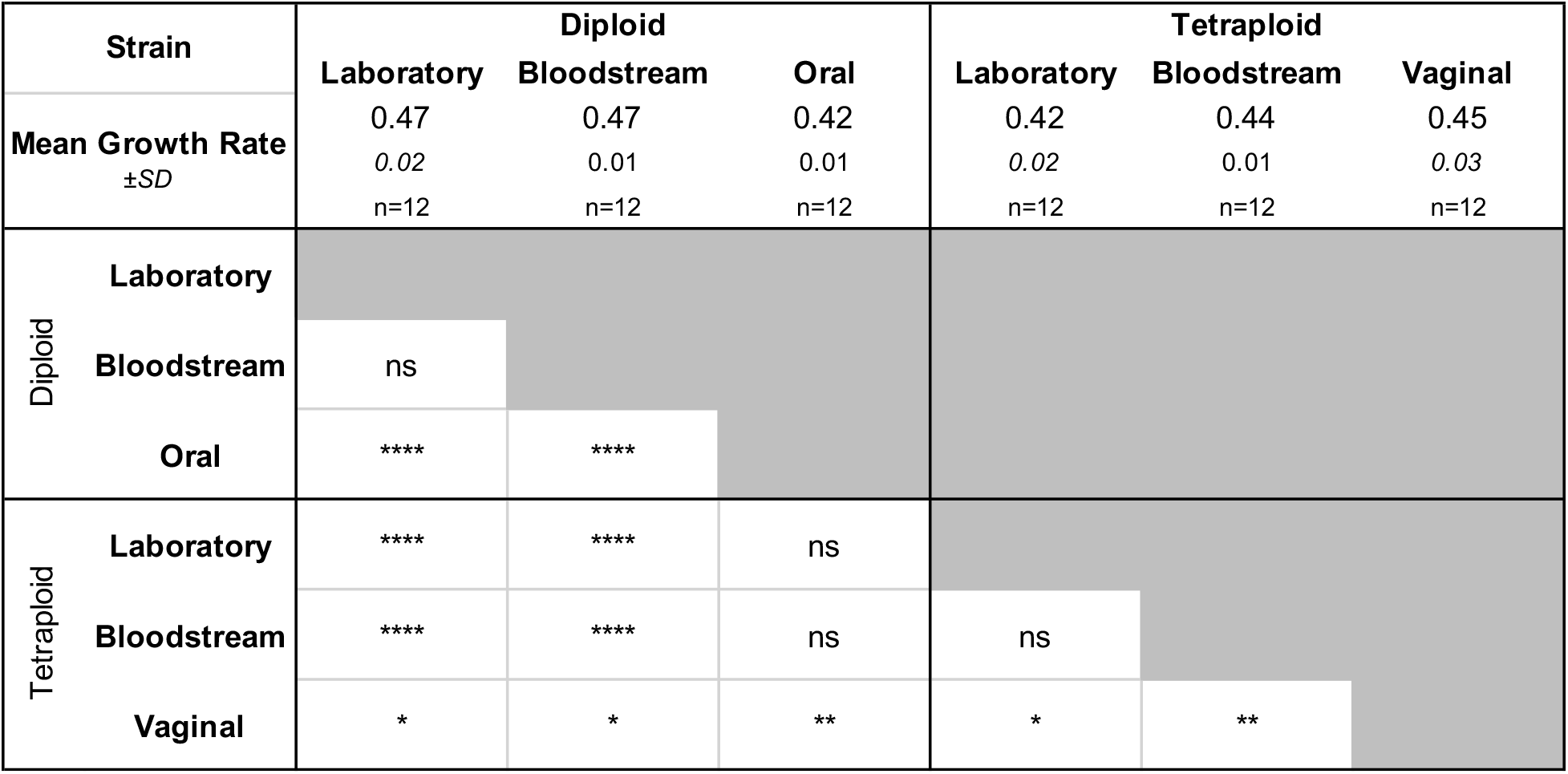
Parental Strain Growth Rates. Mean growth rate, standard deviation (SD) and number of biological replicates (n) are indicated for each treatment. Differences between treatments were tested in all pairwise combinations using the Mann Whitney U test and significance is indicated (ns – not significant; * p<0.05; ** p>0.01; *** p>0.001; *\*\*\*\** p>0.0001).

**Supplemental Table S6:**
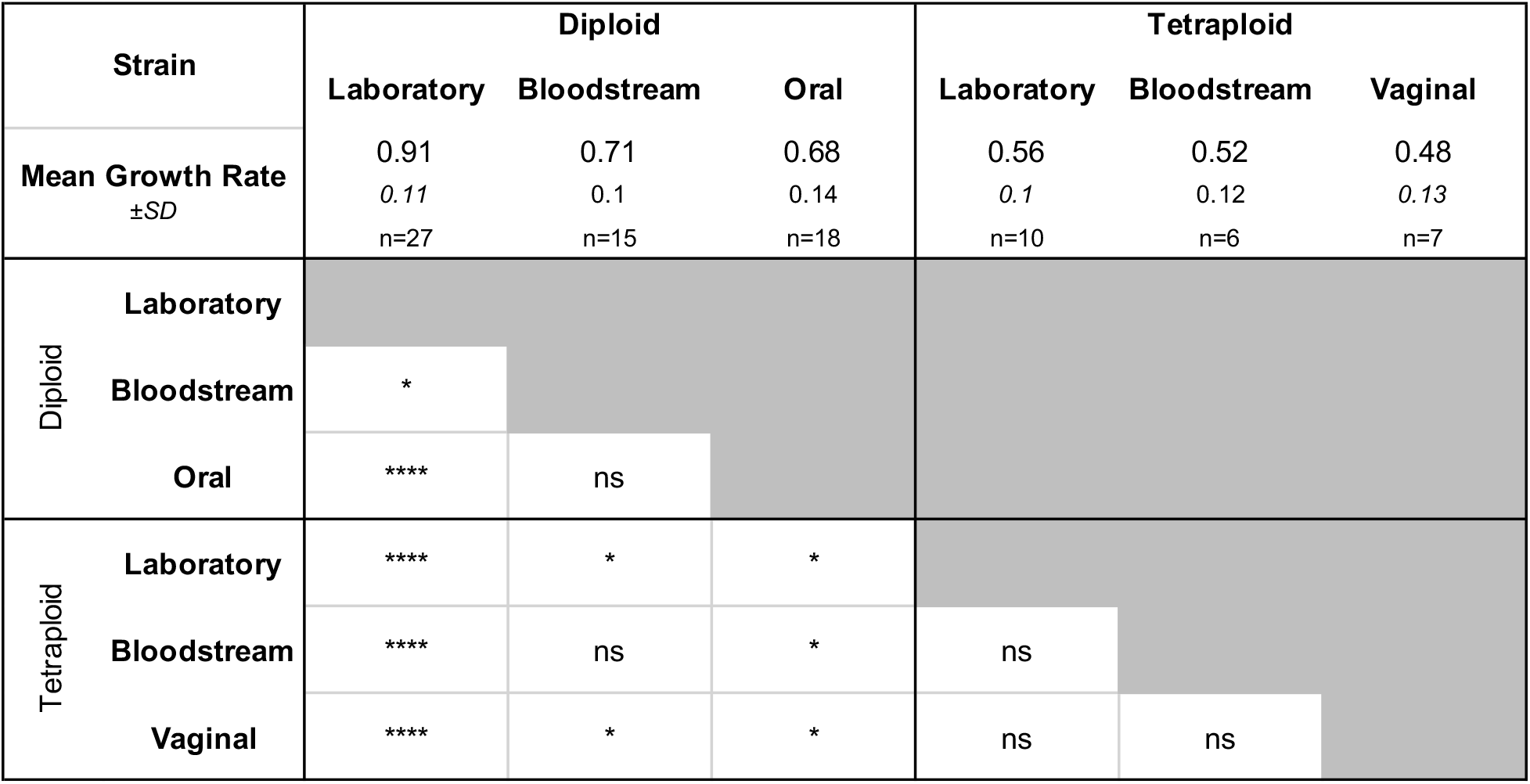
Parental Strain Relative Host Infected Fecundity. Relative fecundity (infected brood size/uninfected brood size), standard deviation (SD) and number of biological replicates (n) are indicated for each treatment. Differences between treatments were tested in all pairwise combinations using the Mann Whitney U test and significance is indicated (ns – not significant; * p<0.05; ** p>0.01; *** p>0.001; **** p>0.0001).

## Notes

### Competing Interest Statement

The authors have declared no competing interest.

